# Multisensory perceptual decision-making persists as alertness fades, even in the absence of an overt response

**DOI:** 10.64898/2026.09.28.754917

**Authors:** Clara Alameda, Daniel Sanabria, Tristan A. Bekinschtein, Luis F. Ciria

## Abstract

Momentary declines in alertness are common in everyday life, yet we must continue to make rapid and accurate decisions from multiple sensory signals that may converge or conflict. Here we investigated multisensory perceptual decision-making across the wake-sleep transition. Participants performed an audio–tactile spatial localisation task while electroencephalography tracked spontaneous fluctuations in alertness. Although drowsiness slowed responses and increased omissions, participants remained largely accurate, and the behavioural influence of congruent and conflicting tactile information persisted. Behavioural modelling revealed slower evidence accumulation and increased decision boundaries. Stimulus- and response-related neural patterns remained robustly decodable across alertness states, despite changes in their temporal dynamics but multisensory congruency was no longer reliably decoded during drowsiness. Brain signatures of lateralized motor preparation persisted during behavioural omissions and, under multisensory conflict, might even favour the competing response. Declining alertness does not uniformly degrade multisensory perceptual decision-making but selectively alters the processes linking sensory information to action.

## INTRODUCTION

Imagine walking through a crowded street late in the evening. Someone calls your name from somewhere behind while, almost simultaneously, you feel a tap on your left shoulder. Both signals immediately interact in your brain, resulting in you turning towards the left to see who called and touched you. This ability to use sensory evidence to guide a choice and an appropriate action lies at the core of perceptual decision-making. Its neural and computational foundations have been extensively characterised in humans and non-human primates during wakefulness, particularly in discrimination and categorisation tasks (Heekeren et al., 2008). Yet perceptual decisions must also be sustained as the brain moves away from optimal wakefulness. How does the brain continue to link sensory evidence to action as alertness fades?

Natural fluctuations in alertness occur continuously throughout the day but become particularly pronounced during the transition from wakefulness to sleep. Sleep onset unfolds as a gradual, dynamic and non-linear process, characterised by transient fluctuations between attentive wakefulness and sleep-like states (Nir & de Lecea, 2023; Ogilvie & Wilkinson, 1984). This transitional period provides a unique experimental model for examining how the processes linking sensory evidence to behaviour are modulated by decreased alertness (Goupil & Bekinschtein, 2012). Across a wide range of cognitive tasks, successful performance can be maintained during the wake-sleep transition despite slower responses, increased errors and intermittent periods of unresponsiveness (Lacaux et al., 2024; Xu et al., 2025). This resilience seems to rely on adaptive changes in the neural and computational processes supporting behaviour, including slower evidence accumulation (Alameda et al., 2024), greater decision volatility (Ciria et al., 2021), emerging spatial biases (Bareham et al., 2014; Jagannathan et al., 2022) and changes in the neural dynamics typically observed during wakefulness (Canales-Johnson et al., 2020).

Most accounts of perceptual decision-making during the wake-to-sleep transition derive from unisensory tasks, leaving unresolved how declining alertness affects decisions built from multiple, potentially conflicting sources of evidence. This is critical because natural behaviour requires the brain not only to register sensory events, but to arbitrate between convergent and competing signals in order to select an action. We recently found that the behavioural and computational benefits of redundant audiotactile stimulation persist as alertness fades, despite marked changes in their underlying neural dynamics (Alameda et al., 2026). Whether the brain can still resolve multisensory evidence into a coherent perceptual choice and translate that choice into action as responsiveness progressively deteriorates remains unknown. What if the voice and the tap on your shoulder came from opposite directions, would drowsiness still allow you to turn in the right direction?

To address this question, we conducted a pre-registered study (https://osf.io/3kgjw/files/yqs9k) in which healthy participants drifted naturally from wakefulness to sleep while performing an audio-tactile spatial localisation task. On each trial, participants reported the location of a task-relevant auditory stimulus while ignoring a simultaneous tactile vibration presented either on the same or the opposite side (see Fig. 1). By combining behavioural, drift-diffusion and EEG analyses, we asked whether decreased alertness produces a unitary breakdown of perceptual decision-making or instead dissociates the component processes linking sensory evidence to action. We show that multisensory perceptual decisions persist as alertness declines. Although decision formation becomes less efficient and canonical conflict-related neural signatures become progressively uncoupled from behaviour, perceptual and response representations remain largely preserved, with motor preparation persisting even beyond the loss of overt behaviour.

**Figure 1.**
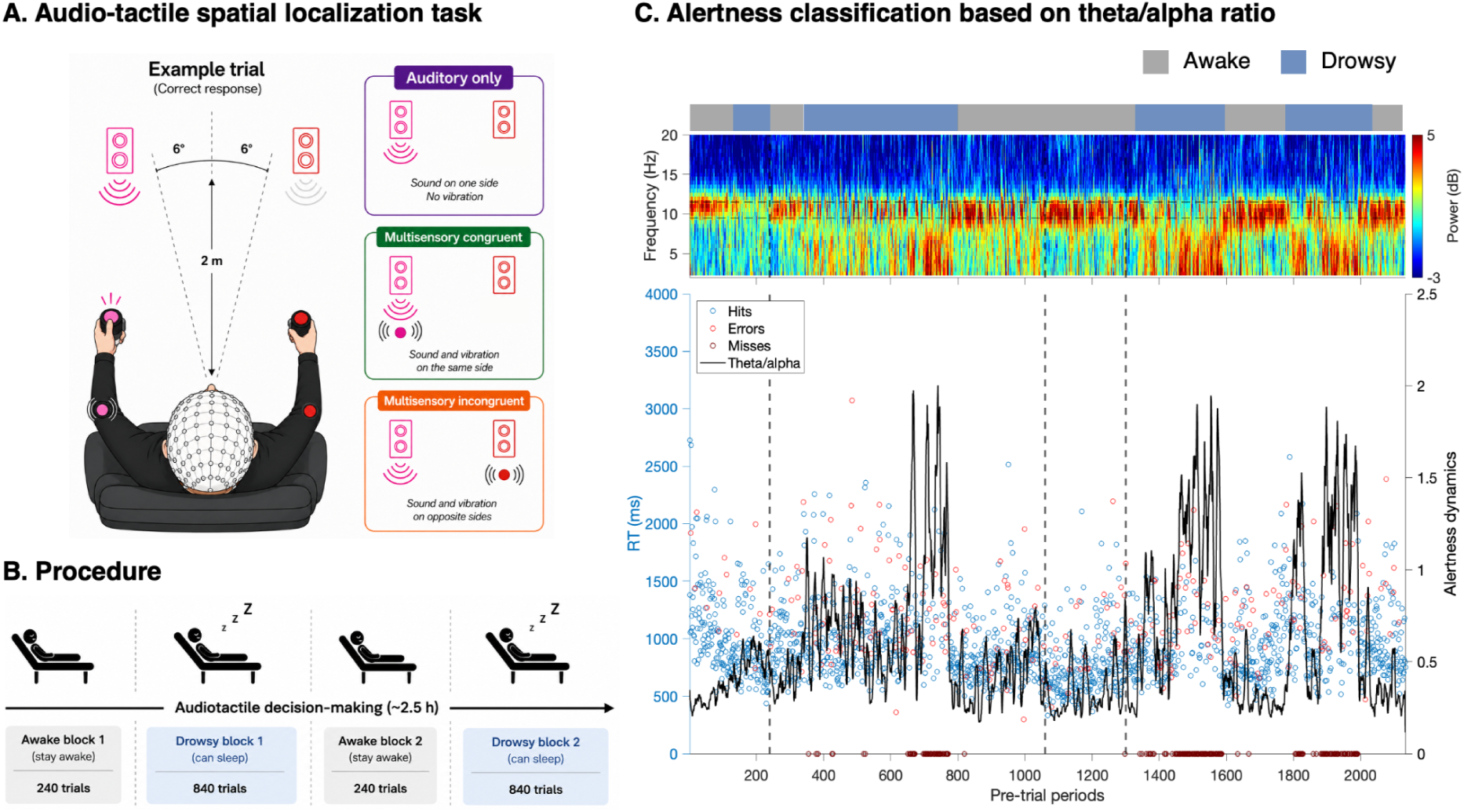
Experimental design and alertness classification. **A)** Right-handed participants discriminated the side (left/right) of a task-relevant auditory stimulus while ignoring task-irrelevant tactile stimulation. Three trial types were presented in random order: auditory-only (left or right), multisensory congruent (same side), and multisensory incongruent (opposite side). Participants were instructed to indicate whether each auditory stimulus was presented to the left or right of the midline (ignoring the tactile stimulus) by pressing the corresponding button with their left or right thumb as quickly and accurately as possible. **B)** The task was divided into four consecutive blocks. During short “awake” blocks, participants were asked to remain alert. During longer “drowsy” blocks, participants were allowed to fall asleep, but if they failed to respond for five consecutive trials, they would be awakened by the experimenter. **C)** Top, time–frequency representation of prestimulus activity (2 s before stimulus onset, occipital electrodes) across the experimental session for one representative participant. Dashed horizontal lines delimit the individual alpha band (IAF ± 1 Hz) used to compute the theta/alpha ratio, while dashed vertical lines indicate the boundaries between blocks. The bar above the spectrogram shows the classification of pre-trial periods as awake (grey) or drowsy (blue). Bottom, trial-by-trial fluctuations in the prestimulus theta/alpha ratio (black line), which was used to classify each trial as awake or drowsy. Coloured circles indicate behavioural outcomes (correct responses, errors and omissions) plotted according to reaction time (0 ms for omissions). Periods of elevated theta/alpha ratio were accompanied by increased reaction-time variability and a higher incidence of omitted responses, consistent with progressive reductions in alertness.

## RESULTS

### Overall performance declines during drowsiness while multisensory interaction effects remain largely preserved

We first asked how reduced alertness affected behavioural performance. As expected, drowsiness produced a marked global deterioration, although participants remained capable of performing the task with relatively high accuracy (∼70% correct responses). Specifically, linear mixed-effects models revealed that participants responded more slowly (*F*(1,330) = 393.95, *p* < 0.001, η²◻ = 0.54), were less accurate (*F*(1,352) = 22.99, *p* < 0.001, η²◻ = 0.06), and failed to respond substantially more often (*F*(1,352) = 493.42, *p* < 0.001, η²◻ = 0.58) than during wakefulness (Fig. 2A-C, left). Participants also showed a small but reliable advantage for right-sided over left-sided stimuli, responding slightly faster (*F*(1,330) = 7.22, *p* = 0.008, η²◻ = 0.02) and more accurately (*F*(1,352) = 30.59, *p* < 0.001, η²◻ = 0.08; Fig. 2A-C, middle). This lateralization was enhanced for accuracy during drowsiness (*F*(1,352) = 4.28, *p* = 0.039, η²◻ = 0.01), consistent with the leftward performance disadvantage previously reported in right-handed participants during sleep-onset (Bareham et al., 2014; Jagannathan et al., 2022).

**Figure 2.**
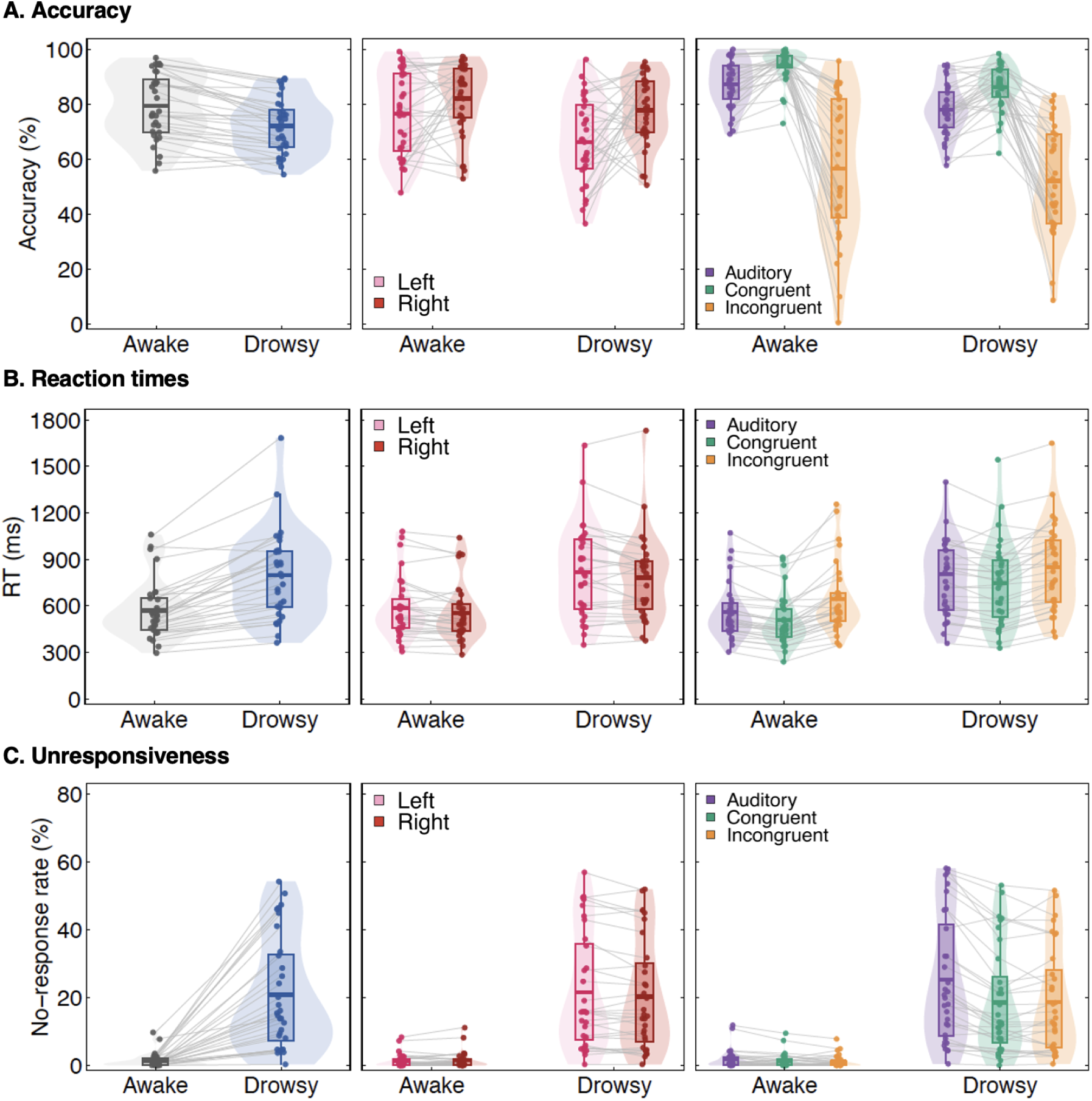
Behavioural performance across alertness states and experimental conditions. **A)** Accuracy (% correct responses), **B)** reaction times (RT) and **C)** omission rate (% trials with no response) are shown as a function of alertness (left column), stimulus side (middle column) and stimulus condition (right column). Reduced alertness was associated with lower accuracy, slower responses and a marked increase in behavioural omissions. Across alertness states, multisensory incongruent stimulation reduced accuracy and increased reaction times relative to auditory-only and congruent trials, whereas both multisensory conditions were associated with fewer omissions than auditory-only trials. Stimulus side exerted only a modest influence on performance, with slightly lower accuracy for left-sided stimuli that became more pronounced during drowsiness. Each dot represents the mean value for one participant in the corresponding condition, and lines connect repeated measurements from the same participant. Boxplots show the median (horizontal line), interquartile range (box) and 1.5× interquartile range (whiskers). Violin plots illustrate the distribution of participant means.

Unlike accuracy and RTs, omission rates were selectively influenced by multisensory stimulation during drowsiness. Both congruent (*t*(357) = 4.46, *p* < 0.001, Cohen’s *d* = 0.75) and incongruent audio-tactile stimulation (*t*(357) = 5.23, *p* < 0.001, *d* = 0.88) significantly reduced omission rates relative to auditory-only trials, indicating that the mere presence of tactile input increased the likelihood of producing an overt response, irrespective of its spatial congruency (Fig. C, right). Despite this overall decline, the behavioural effects of multisensory stimulation were remarkably similar across alertness states. Spatially congruent tactile stimulation facilitated auditory localisation, leading to higher accuracy (*t*(356) = 4.21, *p* = 0.0001, Cohen’s *d* = 0.53) and faster responses (*t*(334) = 3.82, *p* = 0.0005, *d* = 0.50; Fig. 2A-B, right) than auditory-only trials. Conversely, incongruent tactile stimulation produced robust multisensory conflict, reducing accuracy relative to both auditory-only (*t*(356) = 15.17, *p* < 0.001, *d* = 1.91) and congruent trials (*t*(356) = 19.37, *p* < 0.001, *d* = 2.44), while also slowing responses relative to auditory-only (*t*(334) = 4.59, *p* < 0.001, *d* = 0.60) and congruent trials (*t*(334) = 8.41, *p* < 0.001, *d* = 1.09).

We also tested whether these preserved conflict effects translated into adaptive trial-by-trial adjustments in behaviour, but no reliable evidence for conflict adaptation was observed. The interaction between previous- and current-trial congruency did not reach significance for either reaction times (RTs; *F*(1,210) = 3.08, *p* = 0.081, η²◻ = 0.01) or accuracy (*F*(1,210) = 1.41, *p* = 0.237, η²◻ = 0.01), and was not modulated by alertness (RTs: *F*(1,210) = 0.45, *p* = 0.503, η²◻ = 0.002; accuracy: *F*(1,210) = 0.72, *p* = 0.396, η²◻ = 0.003).

### Hierarchical drift–diffusion modelling reveals alertness-dependent changes in evidence accumulation and decisional bias

We then used HDDM to disentangle the factors that modulate decision-making under reduced alertness. We first examined global changes in decision efficiency using an accuracy-coded model, in which the two decision bounds corresponded to correct and incorrect responses (Fig. 3A). Drowsiness was associated with a marked reduction in drift rate relative to wakefulness (Δ = 0.69, 95% CI [0.52, 0.87], *P*(diff > 0) = 1.00), revealing a slower accumulation of sensory evidence towards the correct response. This reduction was accompanied by an increase in boundary separation (Δ = −0.20, 95% CrI [−0.36, −0.05], *P*(diff > 0) = 0.996), confirming that participants required more evidence before committing to a decision. By contrast, non-decision time did not differ reliably between alertness states (Δ = 0.01, 95% CrI [−0.03, 0.05], *P*(diff > 0) = 0.67), suggesting that reduced alertness primarily affected the decision process itself rather than sensory encoding or motor execution (Fig. 3 C-E; full posterior parameter estimates are reported in Supplementary Table S3.).

**Figure 3.**
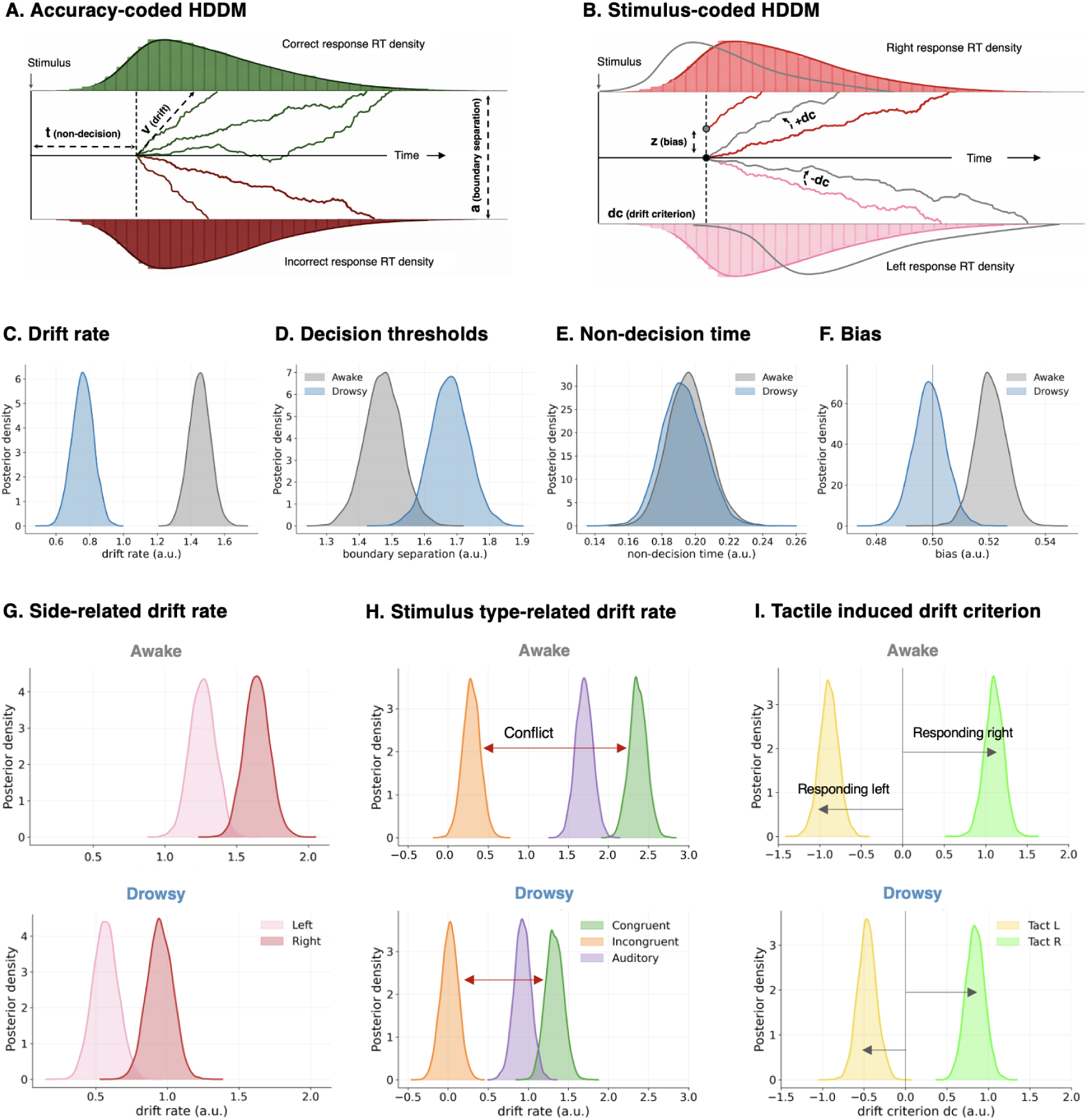
Evidence accumulation models. **A)** Schematic of the accuracy-coded HDDM, in which the decision bounds correspond to correct and incorrect responses. Drift rate (*v*) indexes evidence accumulation, boundary separation (*a*) the amount of evidence required before committing to a decision, and non-decision time (*t*) processes related to sensory encoding and motor execution. **B)** Schematic of the stimulus-coded HDDM, in which left and right responses define the decision bounds. Response bias can be implemented either as a shift in starting point (*z*) or as a drift criterion (*dc*). **C–F)** Posterior densities represent parameter estimates from the final hierarchical models. Drowsiness reduced drift rate (*v*) and increased boundary separation (*a*), whereas non-decision time (*t*) and starting-point bias (*z*) showed little evidence for reliable changes. **G)** Evidence accumulated slightly faster for right-than left-sided auditory stimuli, and this lateralization remained stable across alertness states. **H)** Multisensory congruent stimulation facilitated evidence accumulation relative to auditory-only trials, whereas incongruent stimulation slowed evidence accumulation. Both effects were present in awake and drowsy states but were attenuated during drowsiness. **I)** Task-irrelevant tactile stimulation systematically biased evidence accumulation towards the stimulated side, regardless of the location of the auditory target. This tactile-induced perceptual bias was reliably attenuated during drowsiness.

We next asked whether the behavioural effects of multisensory stimulation were reflected in evidence accumulation. Relative to auditory-only trials, congruent tactile stimulation increased drift rate (Δ = −0.47, 95% CrI [−0.67, −0.26], P(diff > 0) < 0.001), while incongruent tactile stimulation reduced drift rate (Δ = 1.17, 95% CrI [0.97, 1.37], P(diff > 0) = 1.00; Fig. 3H), consistent with slower evidence accumulation under multisensory conflict. Importantly, this conflict-related slowing of evidence accumulation was significantly attenuated during drowsiness (interaction: Δ = −0.60, 95% CrI [−1.03, −0.17], P(diff > 0) = 0.002), revealing that multisensory conflict exerted a weaker influence on evidence accumulation under reduced alertness despite remaining behaviourally robust. A modest advantage was also observed for right-sided stimuli (Δ = −0.14, 95% CrI [−0.24, −0.05], P(diff > 0) = 0.002), suggesting slightly faster evidence accumulation for right-than left-sided stimuli. This lateralisation, however, did not reliably differ between alertness states (interaction: Δ = 0.03, 95% CrI [−0.17, 0.22], P(diff > 0) = 0.61; Fig. 3G)

To examine whether reduced alertness altered decisional or perceptual sources of spatial bias, we also fitted a stimulus-coded model, in which the two decision bounds corresponded to leftward and rightward responses (Fig. 3B). Starting-point estimates (*z*) remained close to the unbiased value of 0.5 in both alertness states, showing only a minimal tendency towards rightward responses that was slightly reduced during drowsiness (Δ = 0.022, 95% CrI [0.006, 0.037], *P*(diff > 0) = 0.997; Fig. 3F). Likewise, drift criterion indicated a modest baseline bias towards rightward evidence accumulation that did not differ reliably between alertness states (Δ = −0.084, 95% CrI [−0.31, 0.14], *P*(diff > 0) = 0.23). In contrast, task-irrelevant tactile stimulation strongly biased evidence accumulation towards the stimulated side (Δ = −1.64, 95% CrI [−1.87, −1.42], *P*(diff > 0) < 0.001). This tactile-driven perceptual bias remained robust in both wakefulness and drowsiness but was attenuated under reduced alertness (interaction: Δ = −0.69, 95% CrI [−1.13, −0.24], *P*(diff > 0) < 0.001). Thus, while task-irrelevant tactile stimulation exerted a strong directional influence on perceptual evidence accumulation, this influence was selectively reduced under lower alertness (Fig. 3I).

### Drowsiness disrupts canonical oscillatory signatures of multisensory conflict

Behaviourally, perceptual decision-making remained largely preserved to reduced alertness, with robust multisensory congruency effects preserved across alertness states. However, HDDM revealed a modest reduction in the influence of multisensory conflict on evidence accumulation during drowsiness. We therefore tested whether the canonical midfrontal theta signature of cognitive conflict (Cavanagh & Frank, 2014) was similarly preserved under reduced alertness.

Consistent with previous work on crossmodal conflict during wakefulness (Morís Fernández et al., 2018), multisensory incongruent stimulation increased midfrontal theta power during wakefulness. Cluster-based permutation tests revealed a significant enhancement of low-frequency theta activity (2–6 Hz, 140–720 ms; *p* = 0.042), consistent with greater engagement of medial frontal conflict-monitoring processes during perceptual decision-making. No corresponding conflict-related cluster emerged during drowsiness. However, the direct contrast between alertness states was not significant, precluding strong evidence that this canonical theta signature was attenuated as alertness declined (Fig. 4A). These findings are consistent with previous observations that canonical theta-band markers of conflict are attenuated across the transition to sleep (Canales-Johnson et al., 2020), despite preserved behavioural conflict effects and only a modest attenuation of the corresponding computational signature, as indexed by evidence accumulation (Fig. 4B).

**Figure 4.**
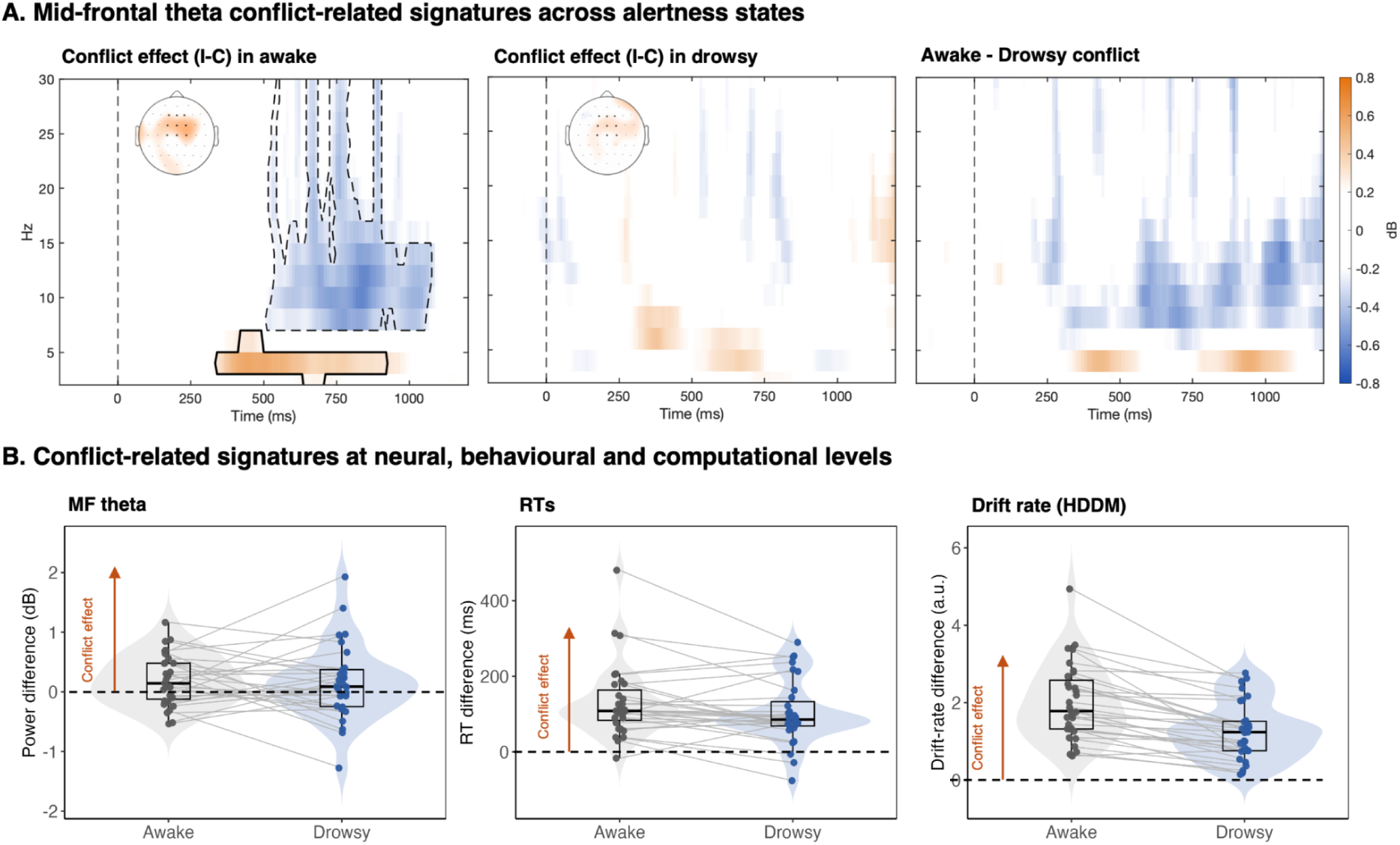
Signatures of multisensory conflict across alertness states. **A)** During wakefulness, multisensory conflict elicited the expected increase in midfrontal theta power (2–6 Hz) over a predefined midfrontal region of interest, whereas no significant theta-band conflict effect was detected during drowsiness. Topographic maps depict the scalp distribution of the awake theta cluster, averaged across its significant time window. **B)** Left, midfrontal theta conflict effect (2–6 Hz) averaged across the significant awake theta cluster. Middle, behavioural conflict effect, quantified as the RT difference between incongruent and congruent trials, showing comparable conflict effects across alertness states. Right, computational conflict effect, quantified as the reduction in HDDM drift rate for incongruent relative to congruent trials, showing a weaker conflict effect during drowsiness. Together, these measures illustrate that while behavioural conflict remains largely preserved during drowsiness, its latent computational signature is attenuated and its canonical midfrontal theta correlate is reduced or even absent. Boxplots indicate the group mean (horizontal line) and interquartile range; individual dots represent participant means, and connecting lines link each participant’s mean across alertness states.

### Multivariate decoding reveals selective preservation of neural representations during drowsiness

Given the loss of a reliable theta-band conflict effect during drowsiness, we used voltage-based multivariate decoding to determine whether task-relevant sensory information, multisensory conflict and response formation remained decodable across successive stages of perceptual decision-making.

#### Stimulus-side representations remain robust during drowsiness

We first examined whether the spatial location of the task-relevant auditory stimulus remained represented during reduced alertness. Decoding was performed on correct-response trials, balancing auditory-only, congruent and incongruent conditions across alertness states. During wakefulness, decoding emerged at approximately 110 ms following stimulus onset and remained reliably above chance throughout the analysed epoch. A highly similar pattern was observed during drowsiness, although decoding emerged slightly later (approximately 190 ms), consistent with a modest delay in the establishment of sensory representations (Fig. 5A, left). Temporal generalization analyses revealed broad within-state generalization in both alertness states. Although neural representations appeared less temporally stable during drowsiness than during wakefulness, robust cross-state decoding was observed from approximately 210 ms onwards, indicating that stimulus-side representations generalized across alertness states (Fig. 5A, middle-right). Thus, task-relevant spatial information remained robustly represented during drowsiness, despite a modest delay in its emergence.

**Figure 5.**
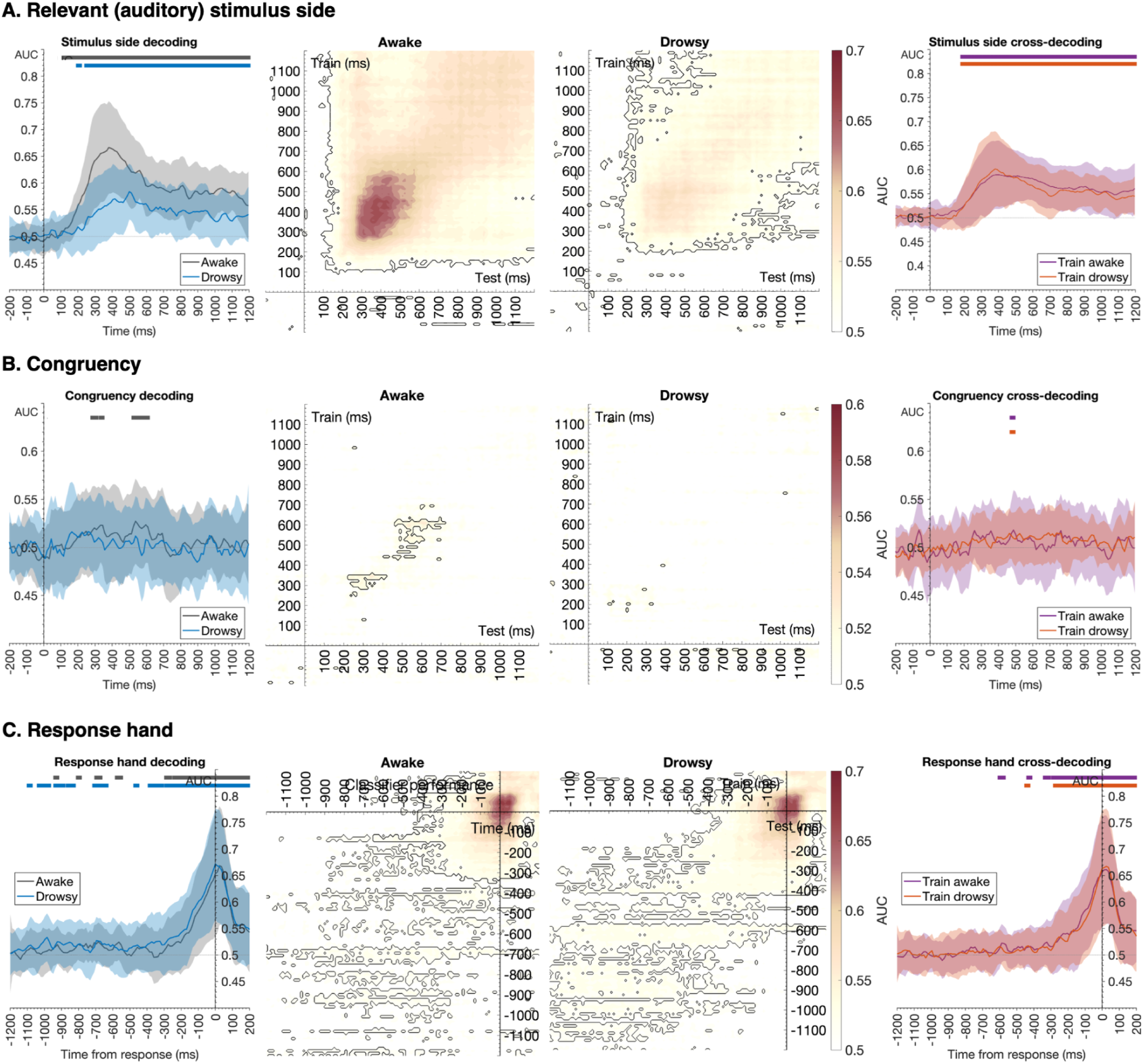
Multivariate decoding of distinct processes supporting perceptual decision-making across alertness states. **A)** Left, time-resolved decoding (AUC) showing robust stimulus-side representations in both alertness states, with a modest delay in decoding onset during drowsiness. Centre, within-state temporal generalization matrices (awake, left; drowsy, right), showing sustained representational dynamics in both states, although representations appeared less stable in time during drowsiness. Right, cross-state temporal generalization showing that stimulus-side representations generalized reliably across alertness states. **B)** Time-resolved decoding revealed only a brief period of above-chance classification during wakefulness, with no reliable decoding during drowsiness. Temporal generalization similarly identified only weak and transient conflict representations during wakefulness, whereas no stable representations emerged during drowsiness. Cross-state decoding suggested that conflict-related neural representations were not generalizable between states. **C)** Response-hand information was robustly decoded in both alertness states before the behavioural response, emerging earlier relative to response execution during drowsiness. Temporal generalization revealed more sustained response-related representations during drowsiness than wakefulness, consistent with slower RT and evidence accumulation. Cross-state decoding remained significant throughout the response preparation period, indicating that motor representations generalized reliably across alertness states despite reduced alertness. Time-resolved decoding (left panels) was performed using voltage-based linear discriminant analysis (LDA) classifiers. Middle panels show within-state temporal generalization matrices (training and testing within awake or drowsy trials), whereas right panels show cross-state temporal generalization (training in one alertness state and testing in the other). Shaded regions indicate ± SEM across participants. Horizontal bars denote significant clusters identified using cluster-based permutation testing.

#### Reduced alertness alters the neural representation of multisensory congruency

We then tested whether multisensory spatial congruency remained decodable in distributed EEG activity. Consistent with the oscillatory analyses, congruency was reliably decoded only during wakefulness, with above chance performance emerging in two post-stimulus windows spanning approximately 270–350 ms and 520–610 ms, whereas decoding remained at chance throughout drowsiness (Fig. 5B, left). Despite the largely preserved behavioural congruency effect, this pattern indicates that the neural representations distinguishing congruent from incongruent multisensory events were no longer robustly expressed under reduced alertness. Consistent with this interpretation, cross-state generalization remained at chance, indicating that the congruency-related patterns expressed during wakefulness were not preserved in a common representational format during drowsiness (Fig. 5B, middle–right). Together with the absence of the canonical midfrontal theta effect, these findings point to a potential state-dependent reconfiguration of the neural processes supporting multisensory conflict, whereby comparable behavioural consequences may be sustained by different neural configurations across alertness states.

#### Response-hand representations remain preserved but become temporally extended

Finally, we examined neural activity associated with response formation and motor preparation by decoding the hand used to execute the behavioural response (left vs. right) from response-locked EEG data. Unlike the previous analyses, response-hand decoding included balanced correct and incorrect trials, thereby reducing the correspondence between auditory stimulus side and response hand and helping to isolate decision- and response-related activity from sensory processing. Response-hand information was robustly decoded in both wakefulness and drowsiness. Sustained decoding emerged approximately 300 ms before the behavioural response during wakefulness and approximately 400 ms before the response during drowsiness, with additional intermittent significant periods extending further back in time under reduced alertness (Fig. 5C, left).

Temporal generalization analyses revealed a corresponding pattern, with response-related representations generalizing over a relatively restricted temporal window during wakefulness. By contrast, these representations extended substantially further backwards in time during drowsiness while maintaining robust cross-state generalization (Fig. 5C, middle-right). Thus, although the neural representations underlying response formation remained largely preserved across alertness states, response-related activity unfolded over a more prolonged time window under reduced alertness. This temporal extension may reflect slower response formation, consistent with evidence accumulation results. Taken together, these findings show that reduced alertness does not uniformly disrupt task-relevant neural representations. Instead, drowsiness preserved sensory and response-related representations while altering the neural representation of multisensory conflict.

### Perceptual decision processes persist even if behavioural responsiveness is lost

To probe whether perceptual decision-making persists beyond overt behavioural responsiveness, we conducted two complementary, non-preregistered exploratory analyses of unresponsive trials during the transition to sleep. Specifically, we assessed whether unresponsive trials simply reflect deeper levels of reduced alertness and, if so, whether perceptual decision-making ceases altogether or whether covert decision processes continue to unfold despite the absence of an overt response.

#### Unresponsive trials occur at progressively deeper levels of reduced alertness

We first compared pre-stimulus theta/alpha ratio across correct responses, incorrect responses and unresponsive trials within the EEG-defined drowsy state. A linear mixed-effects model revealed a robust effect of response outcome (*F*(2, 21441) = 342.90, *p* < 0.001, η²◻ = 0.03; Fig. 6A). Incorrect responses were preceded by higher theta/alpha ratios than correct responses (*z* = −8.18, *p* < 0.001, Cohen’s *d* = 0.14; Fig. 6B), while unresponsive trials exhibited higher ratios than both incorrect responses (*z* = −16.31, *p* < 0.001, *d* = 0.36; Fig. 6C) and correct responses (*z* = −26.17, *p* < 0.001, *d* = 0.50). Therefore, behavioural unresponsiveness occurred during the deepest levels of reduced alertness, with incorrect responses occupying an intermediate stage between successful performance and response failure.

**Figure 6.**
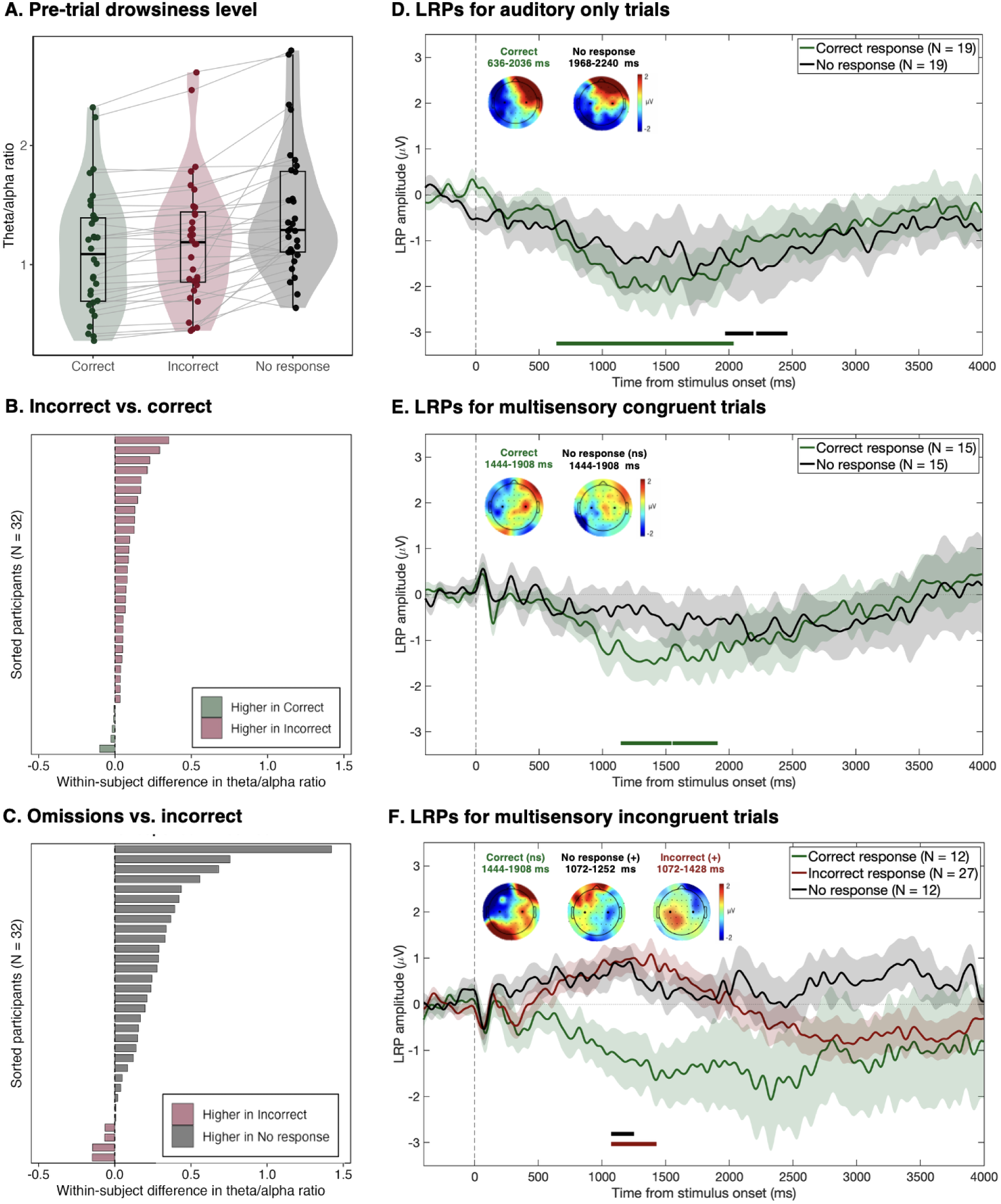
Exploring sleep-related unresponsiveness. **A)** Behavioural performance deteriorated progressively with increasing drowsiness: incorrect responses were preceded by higher theta/alpha ratios than correct responses, whereas behavioural omissions occurred at the deepest levels of reduced alertness. Boxplots indicate the group mean (horizontal line) and interquartile range; individual dots represent participant means. **B)** Subject-by-subject differences in pre-trial theta/alpha ratio between incorrect and correct response, showing higher values for incorrect responses. **C)** Crucially, theta/alpha values were larger for omissions than incorrect responses in most participants. **D–F)** Green traces correspond to correct responses, black traces to behavioural omissions and, in **F**, red traces to incorrect responses. Horizontal bars indicate time intervals during which LRPs differed significantly from zero according to cluster-based permutation testing. Insets show scalp topographies averaged across the corresponding time windows. **D)** In auditory-only trials, both correct responses and behavioural omissions exhibited the canonical negative LRP associated with preparation of the correct response, although the omission-related LRP emerged substantially later. **E)** For congruent trials, correct responses again showed a reliable negative LRP. Omissions exhibited a qualitatively similar waveform and scalp distribution, although the LRP did not differ significantly from zero. **F)** Within incongruent trials, correct-response LRPs did not differ significantly from zero. By contrast, behavioural omissions exhibited a positive LRP closely resembling that observed for incorrect overt responses, consistent with covert preparation of the competing incorrect response rather than the absence of motor preparation. These analyses were exploratory and based on relatively small numbers of participants, and should be interpreted with appropriate caution.

#### Covert motor preparation survives behavioural unresponsiveness

We then examined whether perceptual decision-making continued to unfold despite the absence of an overt response by computing stimulus-locked lateralized readiness potentials (LRPs), an electrophysiological marker of response preparation over premotor cortex (Coles, 1989). Because omissions were relatively infrequent, particularly in the multisensory conditions, many participants did not contribute the minimum number of balanced omission trials required for reliable LRP estimation (30 trials per condition). Consequently, the final samples comprised 19 participants for auditory-only trials, 15 for congruent multisensory trials, and 12 for incongruent multisensory trials.

In auditory-only trials, correct responses exhibited the expected negative LRP, which differed significantly from zero between 636 and 2036 ms after stimulus onset (*p* = 0.002) suggesting that the motor plan associated with the stimulus-appropriate decision might be initiated even in the absence of an overt behavioural response. Importantly, no-response trials also showed a significant negative deviation from zero, although this effect emerged substantially later (1968–2192 ms, *p* = 0.037; 2212–2460 ms, *p* = 0.043; Fig. 6D). The corresponding scalp topographies displayed the canonical lateralisation associated with preparation of the correct response. A similar pattern was observed for multisensory congruent trials. Correct responses again showed a reliable negative LRP that differed significantly from zero between 1144 and 1544 ms (*p* = 0.026) and between 1552 and 1908 ms (*p* = 0.033). Although no-response trials exhibited a comparable negative waveform and scalp distribution, the LRP did not differ significantly from zero at any time point (Fig. 6E). Given the small sample size in this condition (N = 15), this pattern may reflect limited statistical power rather than the absence of covert motor preparation.

Crucially, multisensory incongruent trials exhibited a qualitatively different pattern. Correct-response LRPs exhibited the expected negative morphology, but the waveform did not differ significantly from zero at any time point. Interpretation of this finding should, however, be tempered by the limited sample available for this condition (*N* = 12). By contrast, no-response trials exhibited a sustained positive deviation from zero between 1072 and 1252 ms (*p* = 0.042), closely resembling the positive LRP observed for incorrect overt responses between 1072 and 1428 ms (*p* = 0.019; Fig. 6F). Given that LRP polarity was defined relative to the stimulus-appropriate response, the positive deflection observed in incongruent omission trials is consistent with covert preparation of the competing response rather than an absence of motor preparation. Although exploratory, the overall pattern suggests that behavioural unresponsiveness during drowsiness can coexist with covert motor preparation, whose direction may be shaped by multisensory context.

## DISCUSSION

The present study shows that multimodal information still guides perceptual decisions when alertness declines, even in the absence of overt response. Despite a general deterioration in performance, multisensory influences on behaviour persisted well into drowsiness. Perceptual and response-related information also remained detectable, although their temporal dynamics changed as alertness declined. In contrast, canonical neural signatures of conflict were no longer reliably expressed, despite the preserved behavioural effect of multisensory conflict. Crucially, response preparation persisted even when behavioural responsiveness was lost, but competing tactile information could favour the wrong action.

Drowsiness is known to progressively compromise behavioural performance during the wake-to-sleep transition (for reviews, Andrillon et al., 2024; Lacaux et al., 2024). Our results place part of this decline within the decision process itself, with slower evidence accumulation and increased decision boundaries extending previous findings from unisensory tasks (Alameda et al., 2024; Jagannathan et al., 2022). Against these broad changes, the behavioural influence of multisensory information remained largely stable. Tactile stimuli continued to bias auditory localisation during drowsiness, consistent with their well-established influence on auditory spatial judgements during wakefulness (Bruns et al., 2011; Sanabria et al., 2005). This stability was not necessarily expected as tactile influence could have increased if drowsiness impaired the ability to disregard irrelevant tactile information, or decreased given the selective loss of responsiveness to tactile stimulation recently observed during sleep onset (Alameda et al., 2026). The latter possibility received some support from drift-diffusion modelling, which revealed a weaker tactile-induced perceptual bias during drowsiness, although the difference was modest. This weaker influence on perceptual choice did not translate into reduced responsiveness to multisensory stimulation, since omissions were less frequent than for auditory-only stimuli, suggesting that multimodal inputs may help sustain behavioural responsiveness as alertness declines.

Multisensory behavioral effects remained during drowsiness, but their neural correlates changed when alertness declined. Task-relevant perceptual neural signatures emerged later and became weaker and less temporally stable, whereas those linked to the selected response extended over a longer period before execution, potentially reflecting slower response formation. Conflict-related neural signatures appeared particularly sensitive to declining alertness. During drowsiness, we could no longer reliably detect canonical correlates of conflict, including the increase in mid-frontal theta. This extends previous observations from unimodal conflict (cf. Avancini et al., 2024; Canales-Johnson et al., 2020) to multimodal competition, where such neural dynamics are well established during wakefulness (Morís Fernández et al., 2018; Polley, 2017). This apparent dissociation between behavioral effects and neural correlates suggests that the canonical neural dynamics observed during wakefulness may represent one possible implementation of conflict-sensitive behaviour rather than a necessary substrate. Such an interpretation is consistent with neural degeneracy, whereby similar behavioural outcomes can emerge from different neural configurations across tasks, contexts or brain states (Westlin et al., 2023). However, whether alternative configurations actively support conflict-related behaviour during drowsiness, or whether the relevant signals simply fall below sensitivity of the present analyses remains unresolved.

This brain-behaviour dissociation became even more apparent during periods of unresponsiveness, when response-selective activity persisted despite the absence of overt behaviour. Auditory omissions showed preparation compatible with the instructed response, whereas incongruent multisensory omissions favoured the response indicated by the task-irrelevant tactile stimulus. Previous work has shown that appropriate responses to task-relevant information can still be prepared during semantic categorisation after overt responses cease (Andrillon et al., 2016; Kouider et al., 2014). Our findings extend this preserved capacity to spatial multisensory decisions, while showing that competing evidence can redirect the response being prepared. This stands in marked contrast to periods of preserved responsiveness during drowsiness, in which participants could still resolve the same competition successfully on many trials. Thus, behavioural unresponsiveness may not simply reflect an absence of response preparation. Instead, under multisensory conflict, a competing response may be prepared without ultimately reaching overt execution.

Importantly, these periods of unresponsiveness emerged at the deepest levels of drowsiness sampled here. Pre-stimulus theta-alpha ratios increased progressively from correct responses to errors and were highest before omissions, placing errors and ultimately behavioural unresponsiveness along a continuum of decreasing alertness. This graded relationship closely aligns with evidence that responsiveness during sleep onset depends on the moment-to-moment brain state rather than on discrete sleep stages alone (Comsa et al., 2019; Strauss et al., 2022). Deepening drowsiness may therefore progressively compromise perceptual decisions before behavioural responsiveness is ultimately lost.

Our results challenge the view of declining alertness as a uniform degradation of perceptual decision-making in multisensory contexts. Instead, the transformation of sensory information into action appears to be differentially shaped by momentary downward shifts in alertness, with behavioural outcomes sometimes preserved despite marked changes in the underlying neural dynamics linking perception to action.

## MATERIALS AND METHODS

### Participants

Thirty-five (24 females; 18–30 years) healthy young adults were recruited through local advertisements and the research centre’s participant platform (SONA). Participants were selected to have Epworth Sleepiness Scale scores between 7 and 14 (Johns, 1991). This range reflects a tendency toward increased sleep propensity, typically corresponding to mild levels of daytime sleepiness that do not necessarily indicate the presence of a sleep disorder (Johns, 1992). All had normal hearing, no history of neurological, psychiatric or sleep disorders, and all were right-handed, in order to assess previously reported lateralisation effects (Bareham et al., 2014, 2015). Three participants were excluded due to technical issues: two related to failures in tactile stimulation delivery and one due to excessive trial loss during preprocessing. Thirty-two participants (22 females; 18–30 years) were included in the final analyses. Participants received monetary compensation. The study was approved by the Ethics Committee of the University of Granada (3508/CEIH/2023).

### Experimental task

Participants performed a modified version of the tone localisation task used by Bareham and colleagues (2014). Three stimulus conditions were presented: (a) auditory, (b) spatially aligned audio-tactile, and (c) spatially misaligned audio-tactile (Fig. 1A). These resulted in six configurations: auditory left, auditory right, auditory left–tactile left, auditory right–tactile right, auditory left–tactile right, and auditory right–tactile left. All configurations were presented with equal probability, yielding a total of 2160 trials across the experiment (360 trials per condition), divided into four blocks (awake–drowsy–awake–drowsy; see Procedure). On each trial, participants were instructed to indicate whether the auditory stimulus originated from the left or right loudspeaker by pressing the corresponding button with their thumbs, irrespective of the tactile stimulus. They were asked to respond as quickly and accurately as possible. The inter-stimulus interval varied randomly between 2 and 3 s, and the response window lasted 4 s. No performance feedback was provided. Before the main task, participants completed a 24-trial familiarization block (6 trials per configuration) to confirm detectability and spatial discriminability.

Auditory stimuli consisted of 30 ms white noise bursts (∼65 dB) delivered through one of two loudspeakers positioned ±6° from the midline at a distance of 2.05 m from the participant’s head. Tactile stimuli consisted of 30 ms vibrations (∼100 Hz) delivered via miniature stimulators (18 mm diameter; Dancer Design, UK) attached to each arm (midpoint between shoulder and elbow). Participants wore earplugs (Moldex-Metric, USA) to minimise the audibility of tactile stimulation while ensuring clear perception of auditory stimuli. This was explicitly addressed in the task instructions and verified during a familiarization phase prior to the experiment, during which correct earplug placement and the inaudibility of the tactile stimulation were confirmed. The familiarization was repeated if necessary. In addition, between blocks, we ensured that the earplugs remained properly fitted and comfortable. Stimulus delivery and response recording were controlled using MATLAB (MathWorks Inc., USA) and Psychtoolbox (Kleiner et al., 2007) and delivered via a custom-built audio-tactile response box using hardware-level relays and Arduino-based control. Temporal precision was verified using an oscilloscope, confirming millisecond-level synchronization across auditory, tactile and trigger signals.

### Procedure

Participants attended a single experimental session scheduled either in the morning (around 9:00h) or afternoon (around 14:00h) and were instructed to obtain a normal night’s sleep prior to testing. Upon arrival, they received task instructions and provided written informed consent. Participants were then fitted with a 64-channel high-density EEG system (EASYCAP GmbH, Germany). The experiment was conducted in a comfortable reclining chair, with lights off and eyes closed throughout. The session consisted of alternating alert and drowsy blocks. During alert blocks, participants were instructed to remain awake. Each alert block comprised 240 trials (40 per configuration) and lasted approximately 15 min. During drowsy blocks, participants were allowed to fall asleep. They were informed that the experimenter would wake them if they stopped responding for 5 consecutive trials. To facilitate sleep onset, participants were encouraged to wear comfortable clothing, remove their shoes, and were offered a blanket. Each drowsy block comprised 840 trials (140 per configuration) and lasted approximately 50 min. Following established protocols in our laboratory, the alert–drowsy sequence was repeated twice to maximise the fluctuations in alertness (Fig. 1B). Short breaks (∼5 min) were introduced between blocks to promote re-alerting. During these intervals, lights were turned on, the chair was brought to an upright position, and participants briefly interacted with the experimenter and completed short subjective reports of their ongoing experience. The full session lasted approximately 3.5 h.

### Alertness classification

As the transition from wakefulness to sleep is characterised by a progressive reduction in alpha activity together with an increase in theta activity (De Gennaro et al., 2001), alertness was quantified on a trial-by-trial basis using the EEG theta/alpha power ratio, following previous studies on cognitive and neural changes during drowsiness (Alameda et al., 2026; Bareham et al., 2014; Ciria et al., 2021; Comsa et al., 2019; Xu et al., 2025). Spectral power was computed for each trial in the −2000 to 0 ms pre-stimulus window using a continuous wavelet transform, ranging from 3 cycles at 3 Hz to 8 cycles at 40 Hz. Theta power (4–6 Hz) was averaged across central electrodes (C3, C2). To account for inter-individual variability in alpha frequency, an individual alpha peak frequency (IAF) was identified for each participant within the 8–13 Hz range after removing the aperiodic component of the EEG power spectrum. Alpha power was quantified within an individualized ±1-Hz band centred on each participant’s IAF and averaged across occipital electrodes (O1, Oz, O2). The theta/alpha ratio was subsequently calculated for each trial, yielding a personalized single index of alertness (see Fig. 1C).

For each participant, trials were classified as drowsy or awake based on the theta/alpha ratio distribution. Drowsy trials were defined as the highest 33% of ratio values, and awake trials as the lowest 33%. This threshold was selected in accordance with the preregistered analysis plan and previous studies of alertness fluctuations (Alameda et al., 2026; Bareham et al., 2014; Ciria et al., 2021; Comsa et al., 2019; Xu et al., 2025), with the aim of isolating trials sampling relatively stable neural states at opposite ends of the wake–sleep continuum. Restricting classification to the upper and lower thirds of the theta/alpha distribution preferentially captures periods closer to attentive wakefulness and periods approaching sleep onset, thereby reducing contamination from intermediate alertness states. Following the concept of sleep-state hysteresis (Saper et al., 2010), isolated awake trials embedded within prolonged drowsy periods (≥10 trials) were reclassified as drowsy, and vice versa, to better capture the gradual dynamics of the wake–sleep transition. Although the blocks were designed to facilitate wake–sleep and sleep–wake transitions as many times as possible, final trial classification was determined exclusively by the EEG theta/alpha ratio, irrespective of block. Following trial classification, an average of 666.09 trials (SD = 57.24) per participant were retained in the awake condition and 669.84 trials (SD = 58.18) in the drowsy condition, including correct responses, incorrect responses, and omissions across all stimulus types. The validity of this classification approach was further assessed by inspecting the temporal evolution of the theta/alpha ratio alongside behavioural measures at the single-participant level (see “Alertness_dynamics_plots_subj_by_subj.pdf” in https://osf.io/3kgjw/files/osfstorage).

### Reaction times, accuracy and irresponsiveness

Mean RTs, accuracy and no response rates were computed for each participant and experimental condition. RT analyses included only correct responses and excluded anticipatory responses (<150 ms), errors and omissions. Accuracy was defined as the proportion of correct responses over total responses (range 0–1), and no response rates corresponded to trials without a response over total trials (range 0-1). To ensure task compliance in the presence of incongruent trials, participants were required to achieve at least 60% accuracy in the randomly interspersed auditory-only condition (one third of all trials). All participants met this criterion and were therefore retained for accuracy analyses. For RT analyses, participants with fewer than 5 correct trials in any condition were excluded to ensure reliable estimates and stable within-subject effects in mixed-effects modelling (N = 2; Brysbaert & Stevens, 2018; Miller, 2024).

RTs, accuracy and no response rates were analysed using linear mixed-effects models implemented in the *lme4* package in R (Bates et al., 2014). For each dependent variable, participants were included as a random intercept, while alertness (awake, drowsy), stimulus type (auditory, multisensory congruent, multisensory incongruent), and stimulus side (left, right), as well as their interactions, were initially specified as fixed effects. Model selection followed an information-theoretic approach based on Akaike’s Information Criterion (AIC). Starting from the full model (Alertness × Stimulus type × Side), progressively simpler models were fitted by removing interaction terms and fixed effects down to a null model including only the random intercept. The best-fitting model was identified as that with the lowest AIC (see Supplementary Table S1 for details on AIC values). Statistical inference was performed on the selected model using Type III analyses of variance with Satterthwaite’s approximation of degrees of freedom, as implemented in the lmerTest package (Kuznetsova et al., 2017). Effect sizes are reported as partial eta-squared (η²p). Significant effects were followed up with pairwise comparisons based on estimated marginal means (*emmeans*), with Bonferroni correction for multiple comparisons.

Additionally, to examine whether multisensory conflict elicited sequential adjustments in cognitive control, we analysed behavioural conflict adaptation using linear mixed-effects models. Conflict was defined as the spatial incongruency between task-relevant auditory and task-irrelevant tactile stimulation, requiring participants to resolve competing spatial information while localising the auditory target. RTs and accuracy were modelled as a function of alertness, current-trial congruency, previous-trial congruency and their interactions. Conflict was indexed by the main effect of current congruency, whereas conflict adaptation was assessed by the current × previous congruency interaction. For visualisation, conflict effects were additionally computed as the difference between incongruent and congruent trials, and conflict adaptation was summarized using the four previous-current trial combinations (cC, cI, iC and iI). Participants contributing fewer than 5 trials per condition were excluded from these analyses (N = 1).

### Hierarchical drift-diffusion modelling

To characterise latent decision processes beyond conventional behavioural measures, we applied hierarchical drift–diffusion modelling (HDDM; Wiecki et al., 2013). Drift–diffusion models are sequential-sampling models that describe how decisions are formed in two-alternative forced-choice tasks (Ratcliff et al., 2016). They assume that, when faced with a choice between two alternatives, individuals gradually accumulate noisy evidence favouring one option or the other over time. A decision is reached when the accumulated evidence crosses a decision threshold, triggering the corresponding motor response. In the present task, participants were required to decide whether the target tone originated from the left or right side of space, making auditory localisation a natural two-choice decision problem. By jointly fitting response choices and reaction-time distributions, HDDM makes it possible to infer the latent cognitive processes that give rise to observable behaviour and to determine how these processes are modulated by alertness and stimulus configuration.

To test our preregistered hypotheses, we fitted two complementary families of models targeting distinct aspects of decision-making. First, accuracy-coded models (correct vs. incorrect responses) were used to assess global changes in decision efficiency (see Fig. 3A). These models tested whether alertness, stimulus type (auditory, multisensory congruent, multisensory incongruent), and stimulus side modulated drift rate (*v*), while boundary separation (*a*) and non-decision time (*t*) varied with alertness. Second, stimulus-coded models (left vs. right responses) were used to dissociate perceptual and decisional biases linked to spatial processing (see Fig. 3B). Here, responses were treated as alternative decision bounds, allowing bias to be expressed as a shift in starting point (*z*; decisional bias) or as a drift criterion (*dc*; perceptual bias), enabling us to test whether spatial tactile information induced directional biases and whether these were modulated by alertness.

Within each model family, we followed a hypothesis-driven model comparison approach. For accuracy-coded models, a baseline specification allowed *v* to vary with alertness, stimulus type, and side, while *a* and *t* varied with alertness; reduced and extended variants tested lateralised and non-decision effects. For stimulus-coded models, competing specifications tested whether bias was better captured by *z* or *dc*, and whether these varied with alertness alone or in interaction with tactile direction, following established approaches (Jagannathan et al., 2022; Leite & Ratcliff, 2011; White & Poldrack, 2014). Models were estimated using Markov chain Monte Carlo (MCMC) sampling and included a fixed outlier component (p_outlier = 0.05). Models were compared using the Deviance Information Criterion (DIC), and the best-fitting model (lowest DIC) was retained (Spiegelhalter et al., 2002; Wiecki et al., 2013; see Supplementary Table S2 for DIC values). Model selection used short chains (2,000 samples, 200 burn-in), followed by longer runs (10,000 samples, 1,000 burn-in) for final estimation of the winning models. The best-fitting accuracy-coded model allowed drift rate to vary as a function of alertness, stimulus type, and stimulus side, while boundary separation and non-decision time varied with alertness only. The winning stimulus-coded model implemented bias through starting point, which varied as function of alertness, and drift criterion, which varied as a function of alertness and tactile direction, while the remaining parameters were held constant across conditions.

Statistical inference was based on posterior distributions. For each contrast, posterior samples were subtracted between conditions to obtain a distribution of differences. Effects were considered reliable when the 95% credible interval excluded zero and the posterior probability in the predicted direction was high (*P*(diff > 0) > 0.95 or < 0.05). Posterior overlap was additionally reported as a descriptive index. Convergence was assessed using the Gelman–Rubin statistic (*R^*) across five independent chains (≈ 0.99), indicating satisfactory convergence (Gelman et al., 2003). Reliability was further supported by low Monte Carlo error (< 5% of the posterior SD). Finally, posterior distributions and trace plots were inspected to confirm stable sampling behaviour. The complete HDDM analysis pipeline, including the Jupyter notebook, input data, model outputs, posterior samples and convergence diagnostics is available in the “HDDM” folder at OSF (https://osf.io/3kgjw/files/osfstorage).

### EEG acquisition and preprocessing

EEG data were recorded using a 64-channel actiCHamp system (Brain Products GmbH, Germany) and processed using custom MATLAB scripts together with EEGLAB (Delorme & Makeig, 2004) and FieldTrip (Oostenveld et al., 2011). Data were sampled at 1000 Hz, referenced online to FCz according to the extended 10–20 system, and electrode impedances were kept below 10 kΩ. Continuous data were band-pass filtered between 0.1 and 40 Hz and notch-filtered to remove line noise, then downsampled to 250 Hz. Independent component analysis (ICA) was applied to remove components related to eye movements, blinks, and other noise sources. Bad channels were manually identified and interpolated. Data were epoched relative to stimulus onset from −1.5 to 2 s for stimulus-locked analyses, and from −2 to 1.5 s relative to the response for response-locked analyses. Additional artefact rejection was performed automatically by excluding epochs exceeding ±200 µV (Canales-Johnson et al., 2020). Finally, data were re-referenced to the average across all channels.

### EEG analyses

#### Univariate time-frequency analysis of conflict-related mid-frontal theta

Time–frequency analyses were conducted to characterise neural markers of conflict processing under reduced alertness. Conflict was operationalised as arising from spatial incongruency between task-relevant auditory and task-irrelevant tactile inputs, reflecting crossmodal conflict between competing sensory signals and inducing interference during perceptual decision-making, as described in multisensory integration frameworks (Morís Fernández et al., 2018). Trials were grouped according to alertness (awake, drowsy) and congruency (congruent, incongruent), resulting in four conditions: congruent-awake, incongruent-awake, congruent-drowsy, and incongruent-drowsy. Only correctly responded trials were included in these analyses. Trials were collapsed across the stimulus side to increase statistical power, as the response side was counterbalanced and midfrontal theta is not expected to be lateralised.

Time–frequency decomposition was performed using FieldTrip (Oostenveld et al., 2011), following established procedures (Avancini et al., 2024; Canales-Johnson et al., 2020). Single-trial power was estimated using Morlet wavelets with a constant width of 3 cycles across frequencies. The signal was decomposed between 2 and 30 Hz in steps of 2 Hz, and time-resolved estimates were computed from −400 to 1200 ms relative to stimulus onset in steps of 10 ms. Power values were converted to decibel (dB) units and normalised using a prestimulus baseline from −400 to −100 ms. For each participant and condition, power was averaged across trials and then across a predefined midfrontal region of interest (C1, Cz, C2, FC1, FCz, FC2, F1, Fz, F2). At the participant level, the neural conflict effect was computed as the difference between incongruent and congruent trials, separately for awake and drowsy states. To assess the effect of alertness, these conflict maps were directly compared between states using cluster-based permutation tests (Maris & Oostenveld, 2007), implemented with 5,000 Monte Carlo iterations and a cluster-level alpha of 0.05 across time and frequency dimensions (−200 to 1200 ms). To ensure stable estimates, participants with fewer than 20 correct trials in any condition were excluded from the analysis (N = 1).

#### Voltage-based multivariate decoding

To determine whether task-relevant information remained represented in neural activity under reduced alertness and, if so, whether the underlying neural representations exhibited similar spatiotemporal dynamics to those observed during wakefulness, we performed a series of multivariate decoding analyses targeting complementary stages of the perceptual decision-making process. Building on previous decoding approaches used to study perceptual decision-making (Jagannathan et al., 2022) and conflict monitoring (Canales-Johnson et al., 2020) during fluctuations in alertness, we decoded (i) the side of the task-relevant auditory stimulus, (ii) stimulus congruency, and (iii) response hand. These analyses were designed to track neural representations related to perceptual information, multisensory conflict, and response formation and implementation, respectively.

For stimulus-side decoding, only correct-response trials were included and classified according to the side of the task-relevant auditory stimulus (left vs. right). Auditory-only, congruent, and incongruent trials were pooled and equally balanced across stimulus type, stimulus side, and alertness state. For congruency decoding, only correct-response trials were included and classified as congruent or incongruent, with trial numbers balanced across stimulus side and alertness state to isolate conflict-related information from lateralised sensory activity. For response-hand decoding, response-locked EEG data were classified according to response hand (left vs. right) without baseline correction. Because correct responses are intrinsically coupled to stimulus side, both correct and incorrect trials were included and balanced within each condition. Trial numbers were further balanced across response hand, stimulus side, accuracy, and alertness state. To ensure reliable classification performance, participants were required to contribute at least 30 trials per condition (e.g., awake congruent trials). This resulted in final samples of 27 participants for stimulus-side decoding, 28 participants for congruency decoding, and 29 participants for response-hand decoding.

Voltage-based decoding analyses were performed using linear discriminant analysis (LDA) classifiers implemented in MVPAlab (López-García et al., 2022). For stimulus-side and congruency decoding, EEG data were analysed in stimulus-locked epochs spanning −400 to 1200 ms relative to stimulus onset, using the −400 to −100 ms interval for baseline correction. Response-hand decoding was performed on response-locked epochs spanning −1200 to 400 ms relative to the behavioural response and no baseline correction was applied. Analyses used 5-fold cross-validation with balanced trial counts across conditions and normalization (mean/SD) based on the training data. Classification performance was quantified using the Area Under the ROC Curve (AUC). Statistical significance was assessed using non-parametric cluster-based permutation testing against empirical chance distributions obtained by shuffling trial labels within participants (100 iterations). Group-level null distributions were generated from 10⁵ permutations, and significant clusters were identified using a cluster-forming threshold of *p* < 0.05. To examine the extent to which neural representations generalized across alertness states, cross-state temporal generalization analyses were additionally performed by training classifiers in one state and testing them in the other.

#### Exploratory analyses of unresponsive trials

Although the preregistered analyses focused on trials in which participants produced an overt behavioural response, an important unresolved question is whether perceptual decision-making processes remain active when participants fail to respond during drowsiness. To address this question, we conducted two complementary exploratory analyses restricted to omission trials occurring within the EEG-defined drowsy state.

##### Pre-stimulus alertness preceding correct, incorrect and unresponsive trials

First, we examined whether unresponsive trials reflected a deeper level of reduced alertness than drowsy trials in which participants produced either a correct or an incorrect behavioural response. Rather than averaging observations within participants, pre-stimulus theta/alpha ratios were analysed at the single-trial level using linear mixed-effects models because response outcomes were highly unbalanced during drowsiness, particularly for incorrect responses and omissions. The model included response outcome (correct, incorrect, no response) as a fixed effect and participant as a random intercept. Statistical significance of the fixed effect was assessed using Type III ANOVA with Satterthwaite’s approximation of the degrees of freedom, followed by Bonferroni-corrected pairwise comparisons of estimated marginal means. Partial eta-squared (η²p) and Cohen’s d were reported as measures of effect size.

##### Lateralized readiness potentials (LRPs) during unresponsive trials

Second, we investigated whether covert response preparation continued to develop during omission trials or whether failures to respond reflected an absence of motor preparation. Stimulus-locked lateralized readiness potentials (LRPs), a well-established electrophysiological marker of motor preparation over premotor cortex (Coles, 1989) were computed separately for auditory-only, multisensory congruent and multisensory incongruent trials. Analysing these stimulus types separately avoided averaging across conditions known to differ in motor activation dynamics (Eimer, 1995; Gratton et al., 1992).

For omission trials, participants were included independently within each stimulus condition if they contributed at least 15 expected-left and 15 expected-right trials (i.e., 30 matched trials contributing to LRP waveform). This resulted in final samples of 19 participants for auditory-only trials, 15 participants for multisensory congruent trials, and 12 participants for multisensory incongruent trials. This criterion provided a compromise between maintaining adequate ERP signal-to-noise ratio while maximising participant retention in these relatively infrequent conditions (Boudewyn et al., 2018; Clayson et al., 2013). To facilitate direct qualitative comparisons between overt and covert motor preparation, LRPs for correct responses were computed using exactly the same participants included in the corresponding omission analysis for each stimulus condition. In addition, because incongruent trials yielded a sufficient number of error trials, an additional exploratory LRP analysis was performed for incorrect responses within the incongruent condition using the same inclusion criterion (N = 27).

EEG epochs extended from −400 to 4000 ms relative to stimulus onset (the entire response time window) and were baseline-corrected using the −400 to −100 ms interval. Trials were grouped according to the expected (correct) response side (left vs. right). LRPs were computed using the standard contralateral-minus-ipsilateral subtraction over C3 and C4: C4 − C3 for expected-left trials and C3 − C4 for expected-right trials (Coles, 1989). These two lateralization waveforms were then averaged, such that negative amplitudes reflected preparation of the correct response irrespective of response side. Statistical significance was assessed using one-sample cluster-based permutation tests against zero (Maris & Oostenveld, 2007), testing the a priori hypothesis of sustained negative LRP deflections. Cluster significance was evaluated using 5,000 Monte Carlo randomisations (*α* = 0.05). Because omission and incorrect LRPs in the incongruent condition unexpectedly exhibited a sustained positive-going lateralization, an additional exploratory positive-tailed cluster-based analysis was performed. Although statistical inference was restricted to the LRP waveforms derived from C3/C4, ERPs from all scalp electrodes were retained throughout preprocessing to enable the computation of grand-average scalp topographies. Topographical maps were generated by averaging activity across the temporal extent of significant clusters and computing the expected left minus right response. For conditions in which no significant cluster was observed, topographies were additionally extracted using the corresponding time window from the matched response condition cluster to facilitate qualitative comparison of scalp distributions across response outcomes.

## ACKNOWLEDGEMENTS

This research was supported by a predoctoral fellowship from the Spanish Ministry of Science, Innovation and Universities to the first author (FPU21/00388), by project PID2022-142737NA-I00 funded by MCIN/AEI/10.13039/501100011033 and by FEDER, EU, and by grant RYC2023-045268-I funded by MCIN/AEI/10.13039/501100011033 and by ESF+, both awarded to L.F.C., as well as by Core Funds from the CCC-Lab. We thank Peter Macko (University of Granada) for his invaluable technical support in designing and implementing the experimental setup. We thank our colleagues at the CIMCYC and the Paris Brain Institute (ICM) for valuable discussions and insightful feedback on this work.

## AUTHOR CONTRIBUTIONS

Clara Alameda: conceptualization; methodology; software; formal analysis; data curation; writing – original draft; writing – review and editing; visualization. Daniel Sanabria: conceptualization; methodology; writing - original draft; writing – review and editing; supervision. Tristan A. Bekinschtein: conceptualization; methodology; writing – review and editing. Luis F. Ciria: conceptualization; methodology; software; visualization; writing - original draft; writing – review and editing; supervision.

## SUPPLEMENTARY INFORMATION

Supplementary information and high-quality figures are available via the Open Science Framework repository (OSF; https://osf.io/3kgjw/files/osfstorage).

## CONFLICT OF INTEREST STATEMENT

The authors have declared no conflict of interest.

## OPEN PRACTICES STATEMENT

The experimental protocol, hypotheses and analyses plan were pre-registered in OSF before data collection (https://osf.io/3kgjw/files/yqs9k). All deviations from the pre-registered procedures and exploratory analyses are transparently identified in the manuscript. Raw EEG data, organized according to the BIDS standard, are available on Zenodo (https://zenodo.org/records/21620509). Sorted data and the analysis-specific code used in this study are publicly available in the OSF repository (https://osf.io/3kgjw/files/osfstorage).

## Notes

https://zenodo.org/records/21620509

https://osf.io/3kgjw/files/osfstorage

### Competing Interest Statement

The authors have declared no competing interest.

