## Supplementary material for "Multisensory perceptual decision-making persists as alertness fades, even in the absence of an overt response"

### Supplementary information

**Supplementary Table S1. Hierarchical linear mixed-effects model comparison for behavioural measures.** Candidate models included the full factorial model (alertness × stimulus type × stimulus side), all two-way interaction models, a main-effects-only model, single-factor models, and a null model. Model selection was based on Akaike Information Criterion (AIC), with lower values indicating better model fit after penalising model complexity. Bold values indicate the AIC-selected model used for inferential statistics.

| Behavioural measure | Model | AIC |
|---|---|---|
| **Accuracy** | Alertness + Type + Side | **−308.32** |
| | Alertness × Type × Side | −303.51 |
| | Type × Side | −286.94 |
| | Alertness × Type | −278.56 |
| | Type only | −262.45 |
| | Alertness × Side | −35.74 |
| | Side only | −27.51 |
| | Alertness only | −24.16 |
| | Null | −16.22 |
| **Reaction time** | Alertness + Type + Side | **4547.78** |
| | Alertness × Type | 4555.96 |
| | Alertness × Type × Side | 4560.74 |
| | Alertness × Side | 4610.74 |
| | Alertness only | 4612.65 |
| | Type only | 4806.32 |
| | Type × Side | 4809.03 |
| | Side only | 4832.26 |
| | Null | 4833.24 |
| **Omission rate** | Alertness × Type | **2831.74** |
| | Main effects (Alertness + Type + Side) | 2840.45 |
| | Alertness × Type × Side | 2842.19 |
| | Alertness only | 2850.13 |
| | Alertness × Side | 2852.93 |
| | Type only | 3138.86 |
| | Null | 3141.39 |
| | Side only | 3143.20 |
| | Type × Side | 3144.58 |

**Supplementary Table S2. HDDM model comparison.** Models differed in the decision parameters that were allowed to vary as a function of alertness and task variables and were compared using the Deviance Information Criterion (DIC). Model names correspond to the original HDDM specifications used in the analysis scripts ( https://osf.io/3kgjw/files/osfstorage). Parameter specifications indicate which latent decision parameters were allowed to vary across experimental conditions (*v*, drift rate; *a*, boundary separation; *t*, non-decision time; *z*, starting-point bias; *dc*, drift criterion). Winning models (lowest DIC within each model family) are shown in bold.

| Family | Model name | Parameter specification | DIC |
|---|---|---|---|
| Accuracy-coded | **mA0** | *v* ~ Alertness + Type + Side; *a* ~ Alertness; *t* ~ Alertness | **29937.49** |
| Accuracy-coded | mA2 | *v* ~ Alertness + Type + Side; *a* ~ Alertness; *t* ~ Alertness + Side | 31832.75 |
| Accuracy-coded | mA1 | *v* ~ Alertness + Type; *a* ~ Alertness; *t* ~ Alertness | 32033.88 |
| Stimulus-coded | **mS0_full** | *z* ~ Alertness; *dc* ~ Alertness × Tactile Direction | **22800.29** |
| Stimulus-coded | mS_both | *z* ~ Alertness; *dc* ~ Alertness | 42179.78 |
| Stimulus-coded | mS_z | *z* ~ Alertness | 42250.32 |
| Stimulus-coded | mS_dc | *dc* ~ Alertness | 42256.17 |

**Supplementary Table S3. Posterior parameter estimates from the winning HDDM models.** Values correspond to posterior means, standard deviations (SD), 95% credible intervals (CI), posterior medians (50%), and Monte Carlo sampling error (MC error) for all free parameters estimated in the winning accuracy-coded model (*mA0*) and stimulus-coded model (*mS0_full*). Parameter estimates are reported separately for each condition level included in the model specification. In the accuracy-coded model, *v* denotes drift rate, *a* boundary separation, and *t* non-decision time. In the stimulus-coded model, *z* denotes starting-point bias and *dc* drift criterion. These estimates form the basis for the posterior contrasts reported in the main text. Values are provided for completeness; statistical inference was based on posterior difference distributions rather than direct comparison of raw parameter estimates.

| Parameter | Mean | SD | 2.5% CI | 50% CI | 97.5% CI | MC error |
|---|---|---|---|---|---|---|
| *a*(Awake) | 1.474 | 0.056 | 1.361 | 1.474 | 1.584 | 0.0009 |
| *a*(Drowsy) | 1.678 | 0.057 | 1.565 | 1.678 | 1.791 | 0.0009 |
| *t*(Awake) | 0.196 | 0.012 | 0.173 | 0.196 | 0.221 | 0.0002 |
| *t*(Drowsy) | 0.193 | 0.013 | 0.169 | 0.192 | 0.218 | 0.0002 |
| *v*(Awake, Left, Auditory) | 1.526 | 0.152 | 1.226 | 1.527 | 1.825 | 0.0022 |
| *v*(Awake, Left, Congruent) | 2.173 | 0.154 | 1.868 | 2.171 | 2.475 | 0.0022 |
| *v*(Awake, Left, Incongruent) | 0.093 | 0.154 | −0.208 | 0.093 | 0.391 | 0.0024 |
| *v*(Awake, Right, Auditory) | 1.869 | 0.151 | 1.569 | 1.869 | 2.171 | 0.0021 |
| *v*(Awake, Right, Congruent) | 2.563 | 0.153 | 2.261 | 2.561 | 2.868 | 0.0025 |
| *v*(Awake, Right, Incongruent) | 0.491 | 0.152 | 0.187 | 0.491 | 0.781 | 0.0025 |
| *v*(Drowsy, Left, Auditory) | 0.737 | 0.149 | 0.452 | 0.733 | 1.033 | 0.0025 |
| *v*(Drowsy, Left, Congruent) | 1.178 | 0.150 | 0.884 | 1.175 | 1.472 | 0.0023 |
| *v*(Drowsy, Left, Incongruent) | −0.209 | 0.153 | −0.508 | −0.208 | 0.090 | 0.0022 |
| *v*(Drowsy, Right, Auditory) | 1.120 | 0.151 | 0.826 | 1.120 | 1.413 | 0.0022 |
| *v*(Drowsy, Right, Congruent) | 1.477 | 0.155 | 1.179 | 1.474 | 1.793 | 0.0022 |
| *v*(Drowsy, Right, Incongruent) | 0.249 | 0.152 | −0.053 | 0.248 | 0.546 | 0.0022 |
| *z*(Awake) | 0.521 | 0.005 | 0.510 | 0.521 | 0.531 | 0.00009 |
| *z*(Drowsy) | 0.499 | 0.006 | 0.488 | 0.499 | 0.510 | 0.00011 |
| *dc*(Awake, Tactile Left) | −0.889 | 0.114 | −1.119 | −0.890 | −0.664 | 0.00189 |
| *dc*(Awake, Tactile Right) | 1.099 | 0.114 | 0.871 | 1.100 | 1.318 | 0.00167 |
| *dc*(Drowsy, Tactile Left) | −0.461 | 0.114 | −0.685 | −0.462 | −0.229 | 0.00151 |
| *dc*(Drowsy, Tactile Right) | 0.840 | 0.114 | 0.615 | 0.840 | 1.067 | 0.00184 |
